# Anopheles resistance to deltamethrin can be caused by the increased abundance of an enteric Aeromonas taxon

**DOI:** 10.64898/2026.01.23.701251

**Authors:** Luisa Nardini, Renée Zakhia, Jakub Czarnecki, Emma Brito-Fravallo, Corine Geneve, Konstantinos Mavridis, Thierry Fricaux, John Vontas, Kenneth Vernick, Gaëlle Le Goff, Christian Mitri

## Abstract

The enteric bacteriome of *Anopheles* mosquito vector has been linked with its vectorial competence, however, its influence on insecticide resistance is poorly understood. We found that the depletion of the bacterial microbiome in susceptible *Anopheles* strains, resulting from antibiotic treatment, led to greater than 50% insecticide deltamethrin tolerance compared to untreated mosquitoes. Simultaneous inhibition of cytochrome P450 activity reverted the antibiotic-induced tolerance phenotype, indicating that the antibiotic-induced deltamethrin tolerance is P450-dependent. We found that the antibiotic treatment, while suppressing most enteric bacterial taxa, allowed proliferation of a particular antibiotic-tolerant *Aeromonas* taxon, most closely related to *Aeromonas hydrophila*. Increasing the abundance of this taxon in mosquitoes not treated with antibiotics phenocopied the tolerance phenotype, converting deltamethrin-susceptible *Anopheles* to deltamethrin-tolerant mosquitoes. Collectively, these results highlight a mechanistic interplay in *Anopheles* mosquitoes between antibiotic-induced enteric dysbiosis and cytochrome P450-mediated detoxification that promotes insecticide tolerance. This effect could influence mosquito vectorial capacity, especially in Africa, where auto-medication with antibiotics is highly prevalent.

## Introduction

The enteric microbiome plays a significant role in the physiology of the host organism – from mediating healthy digestion, with dysbiosis linked to various diseases in humans, to inhibiting pathogen development in insect vectors of diseases. The microbiome is probably composed of the most diverse phyla of microorganisms and has also been described as a separate organ inside the host body (1, 2).

In mosquitoes, the diversity of the gut microbiome has been studied on both laboratory strains and field mosquitoes (3) (4) (5) (6). The microbiome is largely derived from the water body inhabited by the larval stages and the diversity is highest at this point (7). When insects reach the adult stage, diversity decreases by approximately 50%, and after a blood meal, it substantially decreases again (7). Adult mosquitoes emerge with at least some carryover of bacteria from the larval and pupal stages. In adult female mosquitoes, a blood meal converts the gut from a carbohydrate-rich environment to a protein-rich one, inducing a significant increase in overall bacterial load and a decrease in diversity of species (7). Mosquito colonies from different geographic origins have been shown to harbor similar microbiome composition regardless of their geographical origin (1, 5), and adult microbiomes of semi-field insects versus laboratory strains are comparable with predominating Enterobacteriaceae and Flavobacteriaceae (7).

Studies related to the mosquito enteric bacteriome have mainly been focused on vector competence. For example, it was shown that the presence of the enteric bacterial flora inhibits the development of *Plasmodium* infection (8), an effect largely due to the gram-negative members stimulating the immune deficiency [IMD] pathway and the production of metabolites such as reactive oxygen species (9). In addition, the microbiome strengthens the gut physical barrier by promoting the synthesis of the peritrophic matrix, which may hamper infection progress (10). Field studies have also correlated specific enteric bacteria taxa with mosquito vector competence. For example, in both wild *A. funestus* and *A. gambiae* from Senegal, the phylum Proteobacteria was found predominant in *P. falciparum*-free vectors (11), whereas Enterobacteriaceae has been positively correlated with *Anopheles* vector competence for *Plasmodium* (12). Consistently, the increase of *Enterobacteriaceae* after trypanosome ingestion in *A. coluzzii* positively correlated with their vector competence for *Plasmodium* (6).

Besides its link to vectorial competence (9), the microbiome has also been involved in the metabolism of xenobiotic compounds such as insecticides, an important feature in the context of crop pests and vector-borne disease (13). Alteration of insecticide susceptibility mediated by the enteric microbiota has been investigated in a variety of urban and agricultural pests, such as the German cockroach (*Blatella germanica*) to indoxacarb (14); the oriental fruit fly (*Bactrocera dorsalis*) to trichlorphon (15); the diamondback moth (*Plutella xylostella*) to chlorpyrifos (16); and the mosquito *Anopheles albimanus* to the pyrethroid, deltamethrin (17). Recently, a study on susceptible and resistant *Anopheles arabiensis* laboratory strains revealed that alteration in the microbiome could influence insecticide resistance (18). Most of these studies have focused on insecticide resistance; however, the influence of the enteric bacteriome in insecticide-susceptible mosquitoes has been poorly studied. For example, in the context of mass drug administration against non-communicable diseases and highly prevalent self-medication with antibiotics in Africa (19), the influence of antibiotic exposures on mosquito susceptibility to insecticides has been little investigated. In addition, the mechanisms underlying the interactions between the enteric microbiome and insecticide susceptibility are poorly understood in mosquitoes.

Here, we investigated the influence of enteric bacteriome dysbiosis mediated by antibiotic exposure on insecticide effect in *Anopheles* mosquitoes. We found that insecticide-susceptible *Anopheles* become tolerant to 50% following antibiotic uptake, which concomitantly decreases the abundance of their enteric bacterial flora. Furthermore, we showed that inhibition of the cytochrome P450 activity reverted the insecticide tolerance phenotype induced by the antibiotic treatment. Finally, we isolated and identified a bacterial taxon from antibiotic-treated mosquitoes that correlates with and may explain most deltamethrin-tolerance phenotype. These findings highlight an interplay between specific enteric microbiome taxa and cytochrome P450 activity, as a mechanism underlying the influence of microbiome dysbiosis upon insecticide susceptibility. This emphasizes the need to consider the role of local interventions, such as antibiotic treatment or self-treatment, as a potential factor promoting the residual malaria transmission by mosquitoes that may escape traditional methods of vector control.

## Materials and Methods

### Mosquito strains and maintenance

The colonies used included three strains of *Anopheles coluzzii* Ngousso, *Anopheles coluzzii* 33S, and *A. stephensi*. *A. coluzzii* 33S is an isofemale line generated in 2015 from a female pedigree originated from Burkina Faso. All colonies were reared under standard conditions (26 °C, 12:12 L:D, and 70% relative humidity) by the Centre for the Production and Infection of *Anopheles* (CEPIA) of the Institute Pasteur in Paris, France. Details of each strain are provided in Table 1.

**Table 1:** Details of colony material used for midgut 16S evaluation and subsequent antibiotic treatments, bioassays and *P. berghei* infections.

| Colony name<br>(species) | Origin | Year colonized |
| --- | --- | --- |
| A.c 33 S (*A. coluzzii*) | Burkina Faso | 2014 |
| A.c Ngousso (*A. coluzzii*) | Cameroon | 2006 |
| *A. stephensi*<br>(SDA500) | Pakistan | 1982 |

### Effects of antibiotic treatment on insecticide tolerance status

Antibiotics were administered by addition to a pre-autoclaved sugar-water solution (10%) on which mosquitoes were allowed to feed *ad libitum*. A broad-spectrum bactericidal antibiotic mix (ATB) of penicillin/streptomycin and gentamicin was added to final working concentrations of 62.5/100μg/ml and 15μg/ml, respectively. Susceptible A.c 33S, *A. coluzzii* Ngousso and *A. stephensi* mosquitoes were provided with antibiotics from emergence for 72 hours, then exposed for 1 hour to deltamethrin (0.05%) using the standard World Health Organization (WHO) tube assay (WHO, 2016), and mortality was scored after 24 hours. Antibiotic-mediated suppression of the microbiome was confirmed by qPCR using primers for total 16S, as described above. Three biological replicates were prepared, and for each, approximately 25 mosquitoes were included per replicate. To test whether the effect could only be due to the chemicals within the mix, the same experiment was performed using the bacteriostatic antibiotic, tetracycline (from Fischer Scientific) (10mg/ml).

To simulate a blood meal on a human host treated with antibiotics, a commonly prescribed antibiotic formulation, amoxicillin, purchased from Sigma-Aldrich, was added to human blood (at 0.2mg/ml) (20). Using a standard membrane feeding assay, female A.c Ngousso were allowed to feed on blood containing (or not, for control) amoxicillin. The mosquitoes were exposed to deltamethrin (0.05%) at 24-, 48- and 72-hours post-feeding using WHO bioassays as described above. Reduction of the microbiome was confirmed by qPCR evaluation of total 16S.

### Bacterial DNA extraction and quantification of total 16S

Evaluation of the total 16S was carried out in insecticide-resistant and susceptible mosquitoes, as well as in antibiotic treated mosquitoes to confirm microbiome suppression. For each experiment, 18 midguts were dissected from pre-cleaned mosquitoes (30 second submersion in 70% ethanol followed by two rinses in sterile PBS 1X solution). DNA was extracted using the DNeasy PowerSoil Kit (Qiagen) according to supplier instructions. DNA was quantified and stored at −80°C. Evaluation of 16S was carried out by qPCR using the bacterial primers 16S Forward and Reverse (Sup Table1) and measured relative to mosquito ribosomal protein S7 (*rpS7*) (Sup Table1). Each reaction comprised 1μl template DNA (2-4ng), 10μl Kapa Sybr Fast qPCR Master Mix (2x), 0.4μl forward primer (10μM), 0.4μl reverse primer (10μM), and 8.2μl nuclease-free H_2_O. Amplification was generated by the following cycling conditions: 95°C/10 min, followed by 40 cycles of [95°C/15 sec.; 60°C/1 min]. Three biological replicates were prepared, and fold changes were computed by the delta-delta Ct method (21). Differences in delta Ct distribution across the independent biological replicates between (dsPara) and (dsGFP) samples were statistically tested using the Student t-test.

**Sup Table 1:**
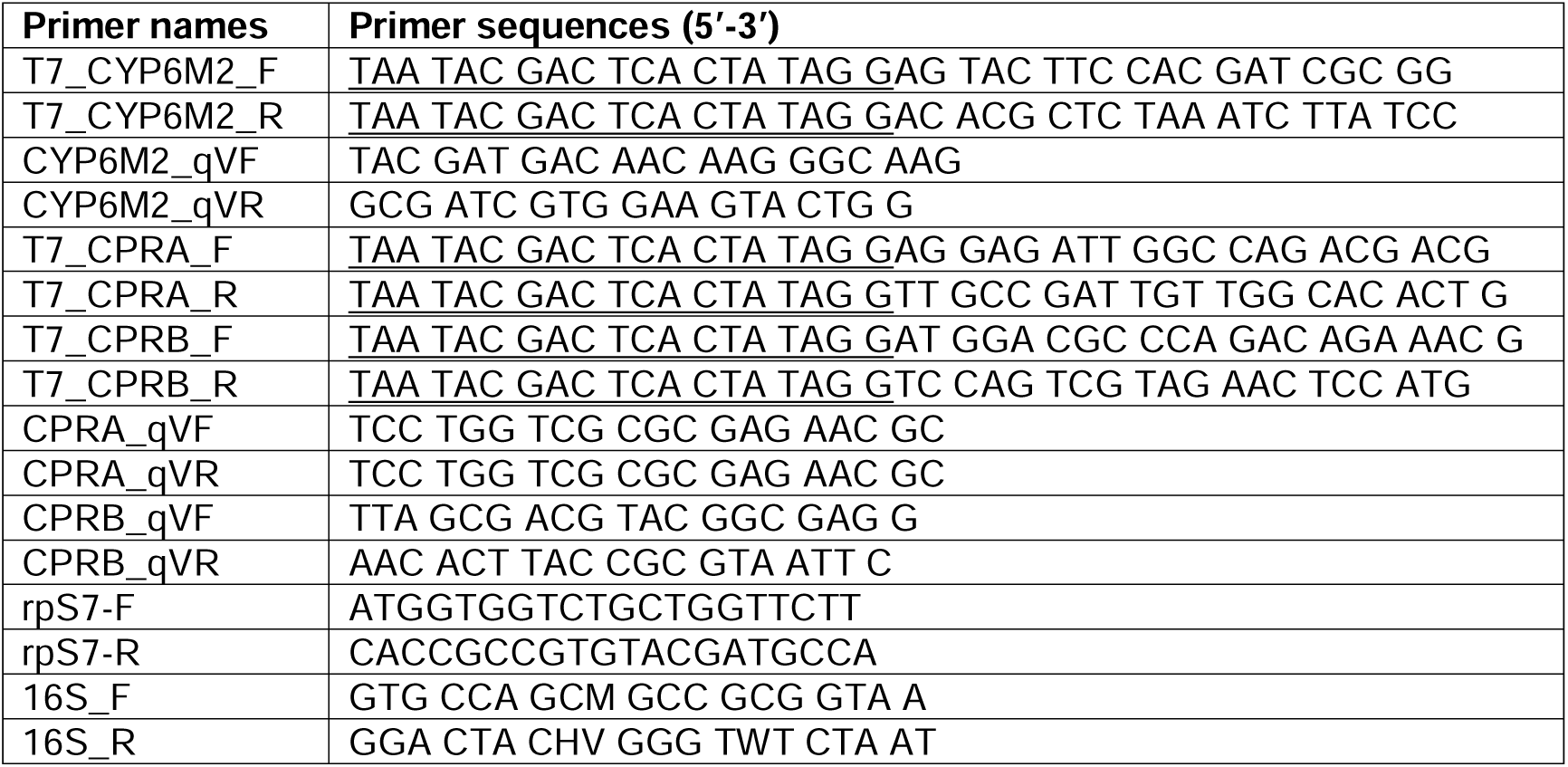
Sequence of Primers used for dsRNA synthesis and for qPCR. Sequence of the primers used for the synthesis of the double-stranded RNA and for knockdown verification. The T7 sequence is underlined for each primer used for dsRNA synthesis. The primers called V are used for the knockdown verification. RpS7 primers were used as an internal calibrator for the qPCR reaction. The 16S primers were used to measure the abundance of the enteric bacteriome.

### Multiplex RT-qPCR for gene expression analysis

The recently described quantitative reverse transcription-real-time PCR (qRT-PCR) 3-plex TaqMan® assay was used for the quantification of the expression levels of eight detoxification genes (CYP6P3, CYP6M2, CYP9K1, CYP6P4, CYP6Z1, CYP6P1 and CYP4G16 and GSTE2) known to play an important role in metabolic detoxification (22), that catalyzes epicuticular hydrocarbon biosynthesis and is implicated in cuticular resistance. Some of these genes were also shown to directly metabolize the pyrethroids (23). The mosquito rpS7 gene was used as an internal calibrator to normalize each reaction for variations in RNA concentrations (22, 24–26). Reactions were performed in the Viia7 Real-Time PCR system (Applied Biosystems) using a one-step RT-PCR mastermix supplied by FTD (Fast-Track Diagnostics, Luxembourg) in a total reaction volume of 10 μL. The thermal cycle parameters were: 50 °C for 15 min, 95 °C for 3 min, and 40 cycles of [95°C/3 sec., 60°C/30 sec.). Samples were amplified in duplicates, and each run always included a non-template control.

### Piperonyl butoxide (PBO) treatment and cytochrome P450 reductase (CPR) silencing

PBO is known to inhibit the activity of P450 (27, 28). PBO was purchased from Sigma-Aldrich and a solution of 4% in olive oil-acetone mixture (1:1) was spotted onto a piece of Whatman (No1) filter paper cut to fit in a standard WHO insecticide assay tube. The paper was dried overnight and placed into a clean bioassay tube to expose the treated surface. Approximately 25 mosquitoes were placed in each bioassay tube and exposed to PBO for 1h, and immediately after, they were exposed to insecticide for one additional hour. Following insecticide exposure, mosquitoes were transferred to a holding tube for 24h when mortality was recorded. A control group of mosquitoes, exposed to PBO only, was included in each experiment.

### Cytochrome P450 reductase and CYP6M2 silencing

To strengthen the role of the P450s in the process, we targeted the P450 reductase, a protein required for electron transfer and activity of all P450s (29). RNAi silencing of the *A. coluzzii* cytochrome P450 reductase (CPR) was performed to reduce the activity of the cytochrome P450s overall. Intrathoracic injections of 500ng/mosquito of dsCPR or dsGFP (control group) were performed on a 1-day-old female A.c 33S treated with the ATB mix from emergence until the insecticide exposure (day 3 days post-injection). Mortality was recorded 24h post-exposure and compared between dsCPR and dsGFP. The primer sequences for dsRNA synthesis and qPCR are available in Sup. table 1. Silencing efficacy was quantified by RT-qPCR as previously described in (30). Briefly, 1 ug of total RNA from dsGFP, dsCPR or dsCYP6M2 was reverse transcribed using random hexamer. Each qPCR reaction comprised 1μl RT product, 10μl Kapa Sybr Fast qPCR Master Mix (2x), 0.4μl forward primer (10μM), 0.4μl reverse primer (10μM), and 8.2μl nuclease-free H_2_O. Amplification was generated by the following cycling conditions: 95°C/10 min, followed by 40 cycles of [95°C/15 sec.; 60°C/1 min]. Three biological replicates were prepared, and fold changes were computed by the delta-delta Ct method (21). Differences in delta Ct distribution across the independent biological replicates between (dsCPR or dsCYP6M2) and (dsGFP) samples were statistically tested using the Student t-test.

### Enteric bacteria culture and transfer

*A. coluzzii* 33S were kept on ATB from emergence for 3 days. At day 4 post-exposure, 20 mosquitoes are dissected to collect their midguts. Midguts were crushed and sonicated (2-3 pulses) in 6mL LB liquid medium incubated at 37°C for 16 hours. At day 5, the bacteria cultured medium was centrifuged at 4000rpm for 5 min to pellet the bacteria and resuspend the pellet into 200μL of PBS. At day 5, a 1:40 dilution of the bacteria resuspended in PBS was performed directly in 2mL of blood and this dilution was used to feed ATB-untreated *A. coluzzii* 33S; a control group was fed on blood without bacteria. Unfed mosquitoes were removed from the cages, and the fed ones were maintained with 10% sucrose at 26°C and 70-80% humidity. 48h post-feeding, mosquitoes fed on blood containing (or not for control) the bacteria were exposed to deltamethrin 0.05%. 24h post-exposure, the mortality was recorded in both mosquito groups.

### Data analysis

Calculation of mRNA gene expression levels in arbitrary units (AU) was calculated using the 2-deltaCt formula. The statistical significances at the *p* = 0.05 level of comparisons were calculated using the two-sided independent samples *t*-test. For bioassay data, mortality rate comparisons of cohort analyses were carried out using the Chi-squared test. The *p*-values of independent tests of significance were combined using the method of Fisher (31). The threshold for significance was set at *p* ≤ 0.01. Statistical analyses were prepared in R (32)

### De novo bacteria sequencing

DNA library was constructed from bacterial DNA from an isolated colony using MGIEasy PCR-Free Library Prep Set (Meghna Group of Industries, China). The sequencing was performed on a DNBSEQ-G400 platform based on a HotMPS High-throughput Sequencing Set. Sequence analyses were conducted using a dedicated pipeline (*de novo*, mapper) available within the Sequana project (33). Kraken 2, a classification system (34), was used to provide and illustrate the taxonomy of the isolated bacteria. This work was performed by the BIOMICS platform at Pasteur Institute.

### Preparation of microsomal fraction

Adult mosquitoes were briefly chilled at –20lJ°C for 1 min and homogenized using a Potter–Elvehjem tissue grinder in homogenization buffer (100lJmM K_2_HPO_4_, pH 7.2, 10lJmg/mL BSA, 1 mM AEBSF, 0.1 mM DTT, 1 mM EDTA). The homogenate was centrifuged at 10lJ000lJg for 5 min at 4lJ°C. A second round of centrifugation under the same conditions was done on the obtained supernatant. Then, the supernatant was centrifuged at 100lJ000lJg for 1 h at 4lJ°C to pellet microsomes, which were washed once with cold phosphate-buffered saline solution and resuspended in microsome buffer (100lJmM K_2_HPO_4_, pH 7.2, 20lJ% glycerol, 1 mM AEBSF, 0.1 mM DTT, 1 mM EDTA). Protein quantification was performed using a Bradford assay with Coo Protein Assay reagent (Uptima, UFP8640) and BSA as a standard curve.

### Cytochrome P450 activity

Microsomal kinetic assays were performed using 7-benzyloxymethoxy-4-(trifluoromethyl)-coumarin (BOMFC) as substrate on three independent mosquito microsome preparations, each tested in three technical replicates over nine substrate concentrations (0.1µM to 200µM). Reactions (final volume 25 µL) contained 5µg protein, (0.1 µM to 200µM) BOMFC in K_2_HPO_4_ 100lJmM pHlJ7.2, and with or without an NADPH-generating system (G6P 2mM, NADP 0.15mM, G6PDH 1U/µL). After 1 h incubation at 30lJ°C, reactions were stopped by adding 25 µL of glycine/NaOH/ethanol stop solution (12.5 mM glycine, 9.7 mM NaOH, 25% ethanol v/v final). Formation of the fluorescent product was measured in a spectrofluorometer (Cary Eclipse, Agilent Technologies, USA) with an excitation and emission wavelength of 390 nm and 497nm, respectively. For each plate, a 7-hydroxy-4-(trifluoromethyl)-coumarin (7-HFC) standard curve (0–25lJpmol/µL) was included to convert fluorescence to product amount. Technical replicates were averaged, and signals without NADPH-generating system were subtracted from those with the system. Corrected values were converted to pmol, initial rates (pmol/min) were calculated and plotted as v=f([S]) for each biological replicate. Apparent Km and Vm values were obtained for each microsomal preparation by nonlinear regression to the Michaelis–Menten equation using the “enzymo” R script (Enzymatic kinetics, 28 October 2006) and reported as mean ± standard deviation of the three biological replicates.

After determining the apparent Km and Vmax values for each condition from kinetic assays, substrate concentrations close to the Km were selected (5 µM for A.c 33S - ATB - BM and A.c 33S – ATB + BM; 10 µM for A.c 33S + ATB – BM, A.c 33S – ATB + BM + C1 bacterial clone and A.c 33S – ATB + BM + C3 bacterial clone) to enable direct comparison of microsomal preparations on the same plate. A two-tailed Student’s t-test was performed to compare A.c 33S - ATB - BM and A.c 33S – ATB + BM, while a one-way ANOVA followed by Tukey’s post hoc test was applied to the three other conditions to assess significant differences among them.

## Results

### Depletion of the enteric bacteria microbiome increases *Anopheles* tolerance to deltamethrin

We tested whether the enteric bacterial microbiome of *Anopheles* susceptible strains could influence deltamethrin toxicity: *A.coluzzii 33S (A.c 33S)*, *A. coluzzii* strain (A.c Ngousso) and *A. stephensi*. Mosquitoes were treated with an antibiotic mix (penicillin, streptomycin and gentamicin) (ATB) from adult emergence for 3 days and exposed to deltamethrin (0.05%). The efficiency of the antibiotic cocktail was confirmed by testing the level of total 16S rRNA (Fig S1). Deltamethrin exposure on ATB-treated mosquitoes rendered both *A.c 33S*, A.c Ngousso and *A. stephensi* tolerant to deltamethrin (0.05%) with 60%, 61%, and 65% mortality respectively in treated groups and 100% in all untreated control groups (A.c 33S: ^2^ = 74.83, df = 10, p = 5.132e-12; A.c Ngousso: ^2^ = 28.91308, df = 6, p = 6.318e-5; *A. stephensi*: χ = 34.43945, df = 6, p = 5.532e-6) (Fig. 1A).

**Figure 1:**
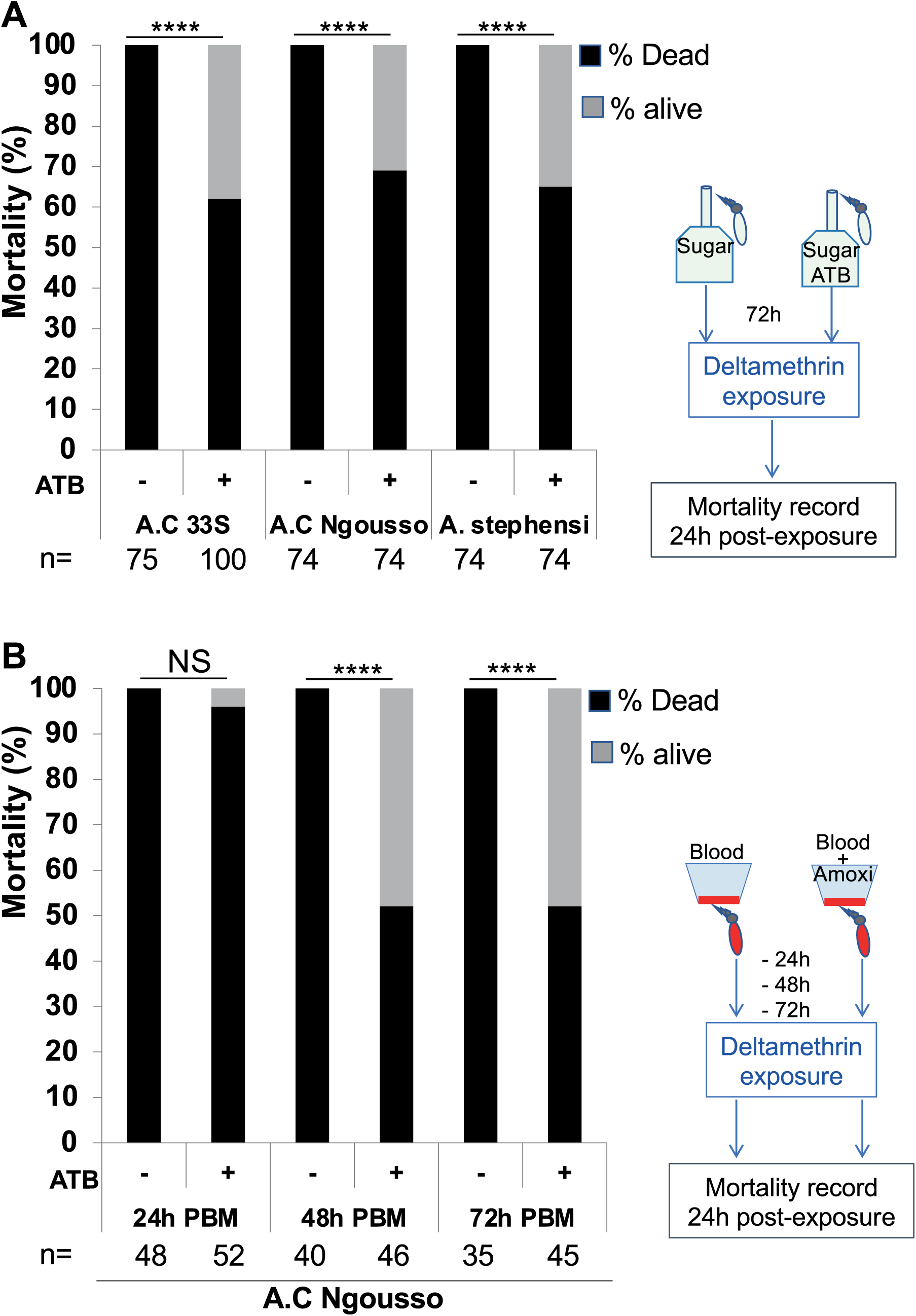
Enteric dysbiosis through antibiotic exposure triggered deltamethrin tolerance in susceptible Anopheles mosquitoes. **A.** Three Anopheles strains (A.c Ngousso, A.C 33S and A. stephensi) were exposed from adult emergence to an antibiotic mixture (ATB) composed of Penicillin/streptomycin and gentamycin for three days before deltamethrin exposure. Mortality rates recorded 24h post-insecticide exposure show that ATB exposure triggered a highly significant tolerance to deltamethrin. This effect was observed in all three tested mosquito strains. ***=p<0.0001**. B.** The histogram shows the influence of a blood meal supplemented with amoxicillin (0.8mg/l) (ATB) compared to a normal blood meal in A. C Ngousso strain. Deltamethrin exposure was performed at 24-, 48 and 72h post-blood meal. Deltamethrin tolerance was observed for the groups exposed at 48 and 72h post-blood meal supplemented with amoxicillin. ***=p<0.0001; n represents the number of tested mosquitoes for each condition.

We then investigate and confirm the correlation between the decreased abundance of the enteric bacteriome and the deltamethrin-tolerance phenotype. Since all three susceptible lines triggered the same phenotype after ATB treatment, we investigated the mechanical link between ATB treatment and insecticide susceptibility in one of them, Ac 33S. ATB-treated A.c 33S survivors from a first insecticide exposure were returned to normal sugar meals (without antibiotic) for 48 hours and re-exposed to deltamethrin (0.05%). Stopping the ATB pressure 48h PBM reverted A.c 33S to their original insecticide susceptible status (∼97% mortality) (Fig S2A). To strengthen the idea that the tolerance phenotype is triggered by enteric dysbiosis and not by any chemical property of the used antibiotics, the same experiment was performed with another antibiotic, Tetracycline, added to the sugar meal at 10mg/ml and acting as a bacteriostatic. This experiment was conducted on two distinct *Anopheles* species, A.c 33S and *A. stephensi* and shows that the tetracycline-exposed group displayed significant tolerance to deltamethrin exposure compared to the unexposed control group (*A.c*: χ^2^ = 5.9801, df = 1, p = 0.01; *A. stephensi*: ^2^ = 6.6576, df = 1, p = 0.009) (Fig S2B). Together with the previously observed phenotype with a different antibiotic mixture (Fig. 1A), this observation under tetracycline exposure indicates that the tolerance phenotype in *Anopheles* is linked to enteric dysbiosis.

We then simulated *Anopheles* mosquito blood-feeding on an antibiotic-treated person to test its influence on insecticide susceptibility. Amoxicillin being a frequently prescribed antibiotic (35–37), A.c 33S were fed on an amoxicillin-supplemented blood meal (0.2mg/ml blood), the control group was fed on blood without Amoxicillin. Mosquitoes were exposed to deltamethrin (0.05%) at 24, 48, and 72 h post-ingestion. Insecticide tolerance was not immediate (i.e., no effect at 24 hours in the amoxicillin-treated group), but at 48 and 72 h post-antibiotic ingestion, we found a statistically significant increase in survival rate in the amoxicillin-treated as compared to control mosquitoes (χ^2^ = 20.2532622, df = 4, p = 0.0004 and χ^2^ = 23.02585, df = 4, p = 0.0001 respectively) (Fig. 1B).

After amoxicillin administration via the blood meal, we observed a progressive reduction of the bacteriome abundance, which reached a maximal at 48h post-blood feeding and rose back at 72h post-feeding (Fig S1B). Antibiotic administration through a blood meal, which by itself enhances the level of the gut flora (3), may explain this profile. The delay between ATB inoculation and the time point required to detect a phenotype suggested either an indirect effect of the ingested antibiotic on insecticide detoxification or that it required 48h for Amoxicillin to reduce the abundance of the enteric bacteriome efficiently (as highlighted in Fig. S1B). Consistently, the tolerance effect was observed from 48h post-amoxicillin ingestion.

### Antibiotic treatment triggering deltamethrin tolerance is P450-dependent

The implication of cytochromes P450 in insecticide detoxification and resistance has been well established (38–40). We tested the role of the P450 activities in the deltamethrin tolerance phenotype mediated by the bacteriome dysbiosis. The synergist piperonyl butoxide (PBO) described as a potent inhibitor of P450 activity, was used to test this hypothesis (27, 28). Sensitive *A. coluzzii* lines were treated with ATB from emergence. After 3 days, they were first exposed (or not for control) to PBO for 1 hour and then to deltamethrin (0.05%) for an additional hour, and mortality was measured 24 hours post-deltamethrin exposure. As described above, the control group (PBO-unexposed) under ATB pressure became tolerant. In contrast, this phenotype was cancelled in the PBO-exposed group under ATB pressure, as 100% mortality was observed in this treatment for both the A.c Ngousso and A.c 33S strains (Fig 2A and 2B). In both lines, the mosquitoes treated with PBO alone (without deltamethrin exposure) were 100% alive (Fig 5), indicating that the mortality observed in the groups exposed to both PBO and deltamethrin is not due to a toxic effect of PBO.

**Figure 2:**
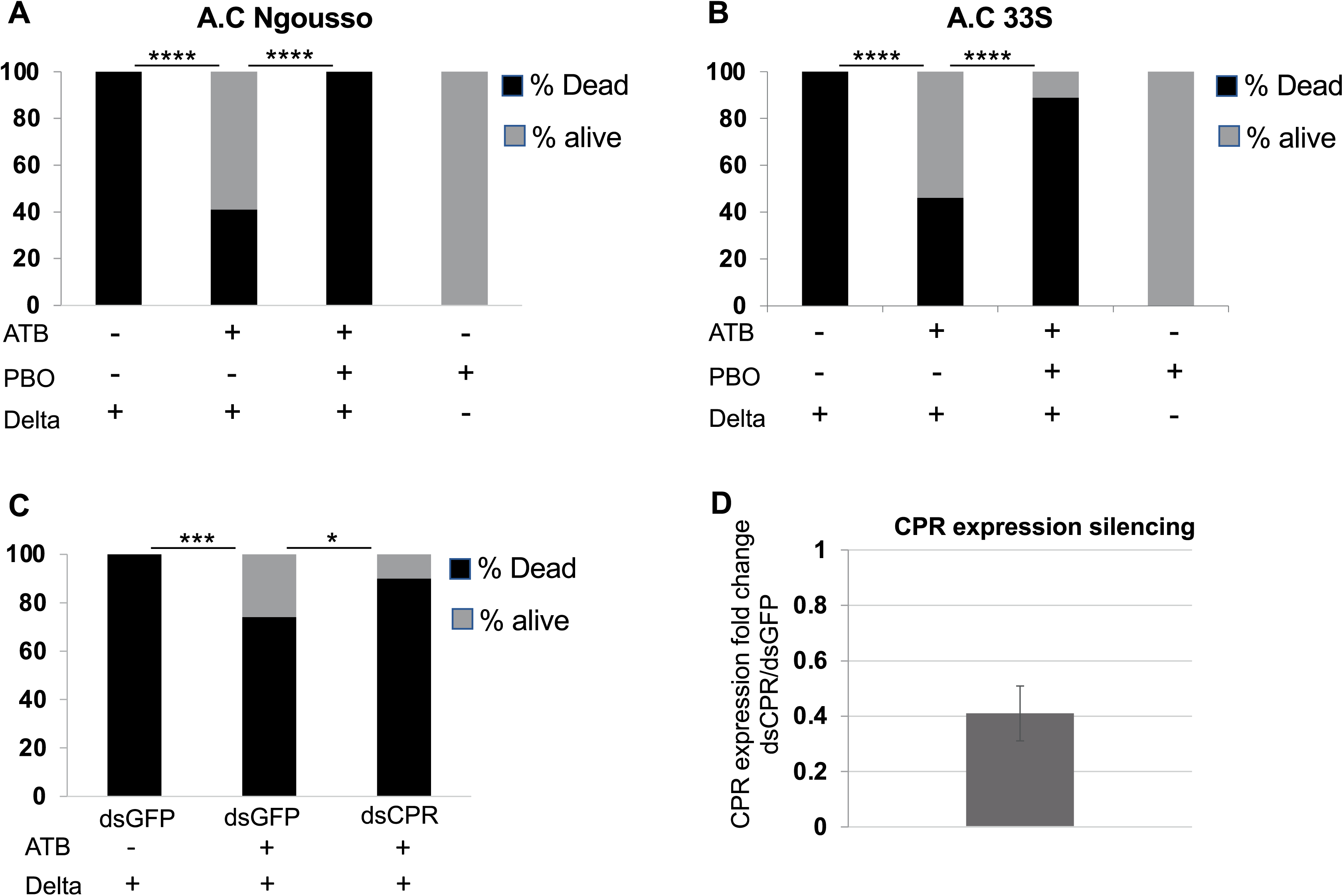
Inhibition of cytochrome P450 abolished deltamethrin tolerance mediated by antibiotic treatment in *A. coluzzii*. The histograms A and B show the influence of pretreating *A. coluzzii* Ngousso (A panel) and *A. coluzzii* 33S (B panel) with the PBO synergist, a P450 inhibitor, on deltamethrin tolerance mediated by antibiotic. We found that deltamethrin tolerance observed in antibiotic (ATB)-treated background was abolished (in A) or significantly reduced (in B) when mosquitoes were pretreated with PBO. The mortality rescue effect is not due to the toxicity of PBO because treatment with PBO alone does not trigger mortality in the tested mosquitoes. C. Silencing of the cytochrome P450 reductase (CPR) enzyme required for P450 activity significantly reduced deltamethrin tolerance mediated by antibiotic treatment. D. The graph shows the silencing efficiency of the CPR transcript by RNAi assays. The expression of the CPR is reduced by 60%. ***=p<0.0001; ***=p<0.001; *=p<0.5; n= total number of mosquitoes. ATB= antibiotics; PBO= pyperonyl butoxide; Delta= deltamethrin.

To strengthen the implication of the P450 activities, we used RNAi-mediated gene silencing to decrease the expression of the cytochrome P450 reductase (CPR), which acts as an electron transporter required for the activity of the P450s (41, 42). ATB-treated A.c 33S were injected with double-stranded RNA (dsRNA) specific for CPR (dsCPR) or control dsRNA for GFP (dsGFP). The two groups were subsequently exposed to deltamethrin three days post-injection, when CPR silencing had reached approximately 60% (Fig. S3A). The recorded mortality showed that CPR expression silencing significantly reduced the tolerance phenotype mediated by the ATB treatment compared to the dsGFP control group (Fig 2B). Together, these results indicate that antibiotic treatment leading to deltamethrin tolerance in *Anopheles* is more likely P450 dependent.

### ATB-treated A.c 33S mosquitoes do not display a clear transcriptomic footprint for known insecticide-detoxifying cytochrome P450s

Our results show that P450 activity is required for the tolerance phenotype mediated by the antibiotic treatment. Therefore, we investigated the transcriptome profile of known P450s, whose function in deltamethrin resistance has been established (22, 24–26). qRT-PCR 3-plex TaqMan® assays were used for the quantification of the expression levels of eight detoxification genes, including P450s (*CYP6P3*, *CYP6M2*, *CYP9K1*, *CYP6P4*, *CYP6Z1*, *CYP6P1, CYP4G16*) and the *GSTE2* between ATB-treated and non-treated A.c 33S. None of the tested genes was significantly modulated in their expression by the ATB treatment. Nevertheless, although not statistically significant, the results showed a slight expression increase of CYP6M2 in ATB-treated mosquitoes as compared to the non-treated ones. CYP6M2 is known to be a direct metabolizer of pyrethroids (43). Therefore, we queried the role of CYP6M2 by RNAi assays in ATB-treated A.c 33S. ATB-treated A.c 33S were injected with dsGFP (as control) or dsCYP6M2 and the injected mosquitoes were exposed to deltamethrin 3 days post-injection. We found no difference in the mortality rate post-exposure between dsGFP and dsCYP6M2 mosquitoes (Fig. 3B), although the expression of CYP6M2 was efficiently silenced (Fig S3). Together, these data suggest that either other detoxifying enzymes or a combinatorial activity of multiple ones may explain the tolerance phenotype observed in the ATB-treated background.

**Figure 3:**
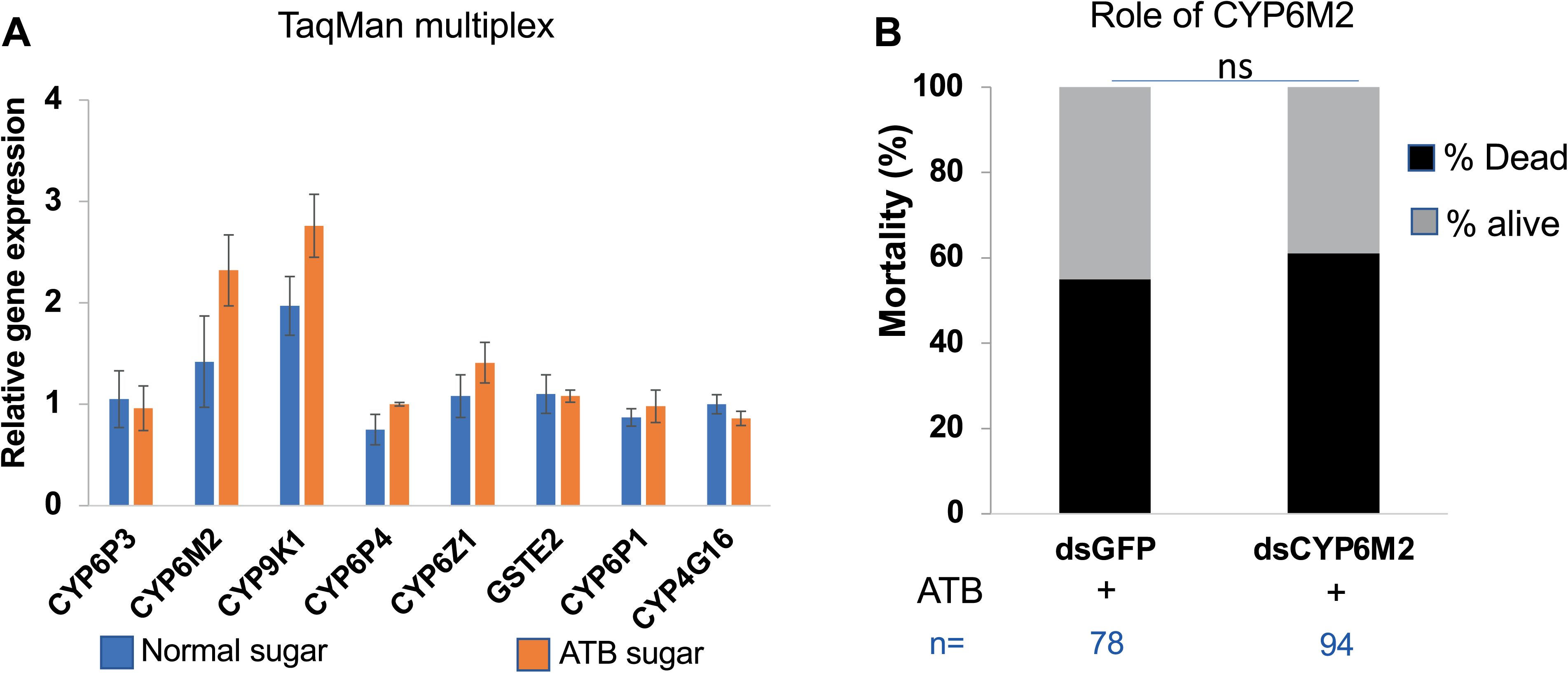
ATB-treated A.c 33S does not trigger a clear transcriptomic footprint on known detoxifying enzymes. **A.** TaqMan assays were performed by targeting 8 detoxifying coding genes, including 7 P450 (CYP6P3, CYP6M2, CYP9K1, CYP6P4, CYP6Z1, CYP6P1, CYP4G16) and one gene coding for the GSTE2. None of the tested genes showed a significant difference in their expression between the ATB-treated group (orange bar) and the untreated control group (blue bar). **B**. Although not statistically significant, a slight increase in expression was observed for CYP6M2. Therefore, we tested the influence of this gene by RNAi. A.C 33S mosquitoes were treated with ATB from adult emergence and injected with dsCYP6M2 and dsGFP. We found that silencing of CYP6M2 did not influence the insecticide tolerance effect mediated by the ATB treatment. n= total number of mosquitoes.

### Aeromonas bacteria OTU isolated from ATB-treated 33S is required for the tolerance phenotype

We hypothesized that expansion of antibiotic-tolerant bacterial taxa in ATB-treated mosquito midguts could induce detoxification enzymes, such as cytochrome P450s, thereby promoting deltamethrin tolerance. To test this, midguts from ATB-treated *A. coluzzii* were collected for bacterial extraction and culture.

*A. coluzzii* females maintained on normal sugar (without ATB) were allowed to feed on blood supplemented or not (for the control group) with the enteric cultured bacteria. 48h after the blood meal, mosquitoes were exposed to deltamethrin, and mortality was recorded 24h post-exposure. This 48h post-blood feeding choice was based on the tolerance phenotype found at the same time point in mosquitoes that had ingested blood supplemented with amoxicillin. We found that mosquitoes fed on the blood supplemented with the cultured enteric bacteria display a tolerant phenotype as compared to the control group or the unfed group (Fig 4A). This result suggested that enteric bacteria from ATB-treated *A. coluzzii* could trigger the tolerance effect.

**Figure 4:**
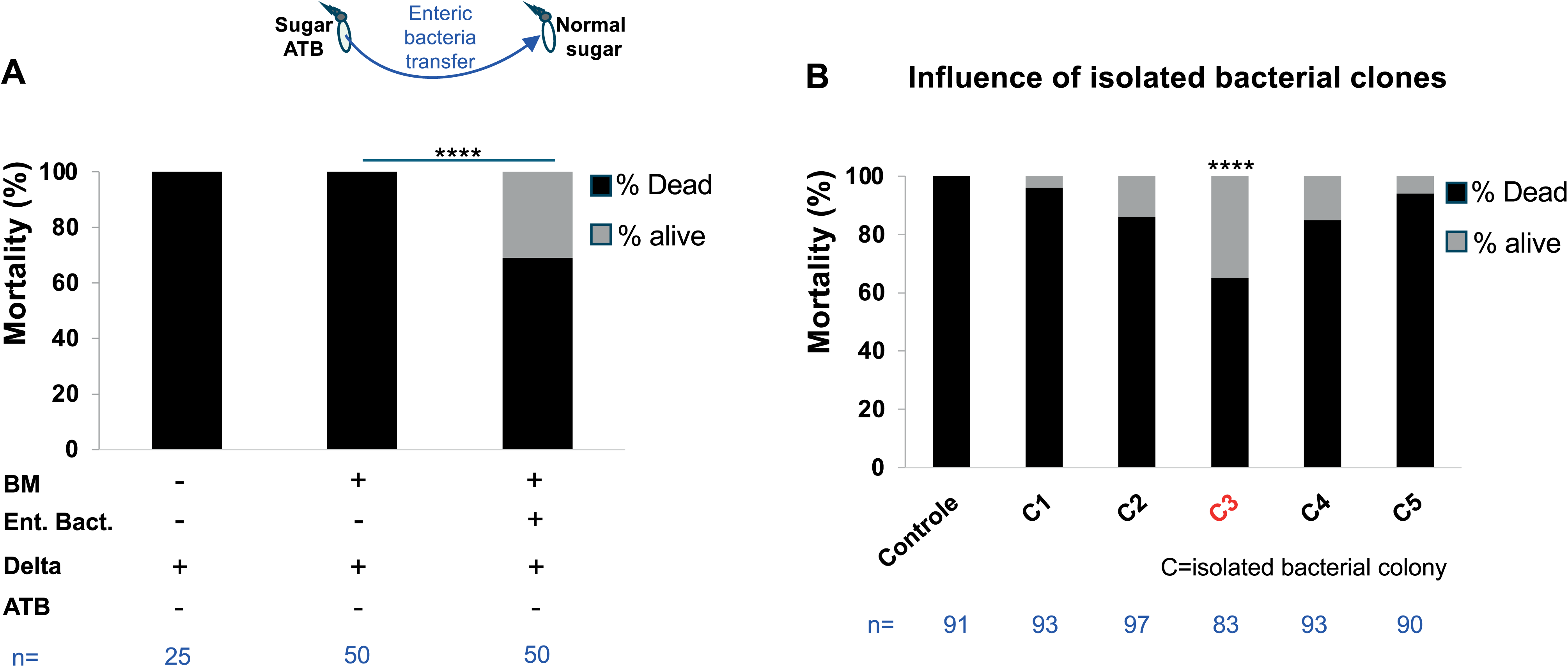
An antibiotic-tolerant taxa belonging to the *Aeromonas* genus is responsible for the deltamethrin tolerance effect. **A.** The graph shows the effect of enteric bacteria transfer from ATB-treated A.C 33S mosquitoes to untreated ones by blood feeding on deltamethrin susceptibility. Deltamethrin exposure was performed 48 hours post-blood meal. The enteric bacteria transfer in untreated (no antibiotic) mosquitoes phenocopied deltamethrin tolerance effect found in the antibiotic-treated background. The naive blood meal did not alter deltamethrin susceptibility compared to unfed mosquitoes. (BM= blood meal; Ent.Bact.= enteric bacteria; Delta= deltamethrin; ATB= antibiotic). **B.** Isolated bacteria colonies from the enteric bacteriome of antibiotic-treated mosquitoes were individually administrated via a blood meal. Mosquitoes were exposed to deltamethrin at 48h post-blood meal. The mosquitoes fed with the bacteria colony N°3 (C3) displayed a significant deltamethrin tolerance compared to the control group. Increasing the abundance of the C3 bacteria taxon can explain deltamethrin tolerance mediated by antibiotic treatment.

**Figure 5:**
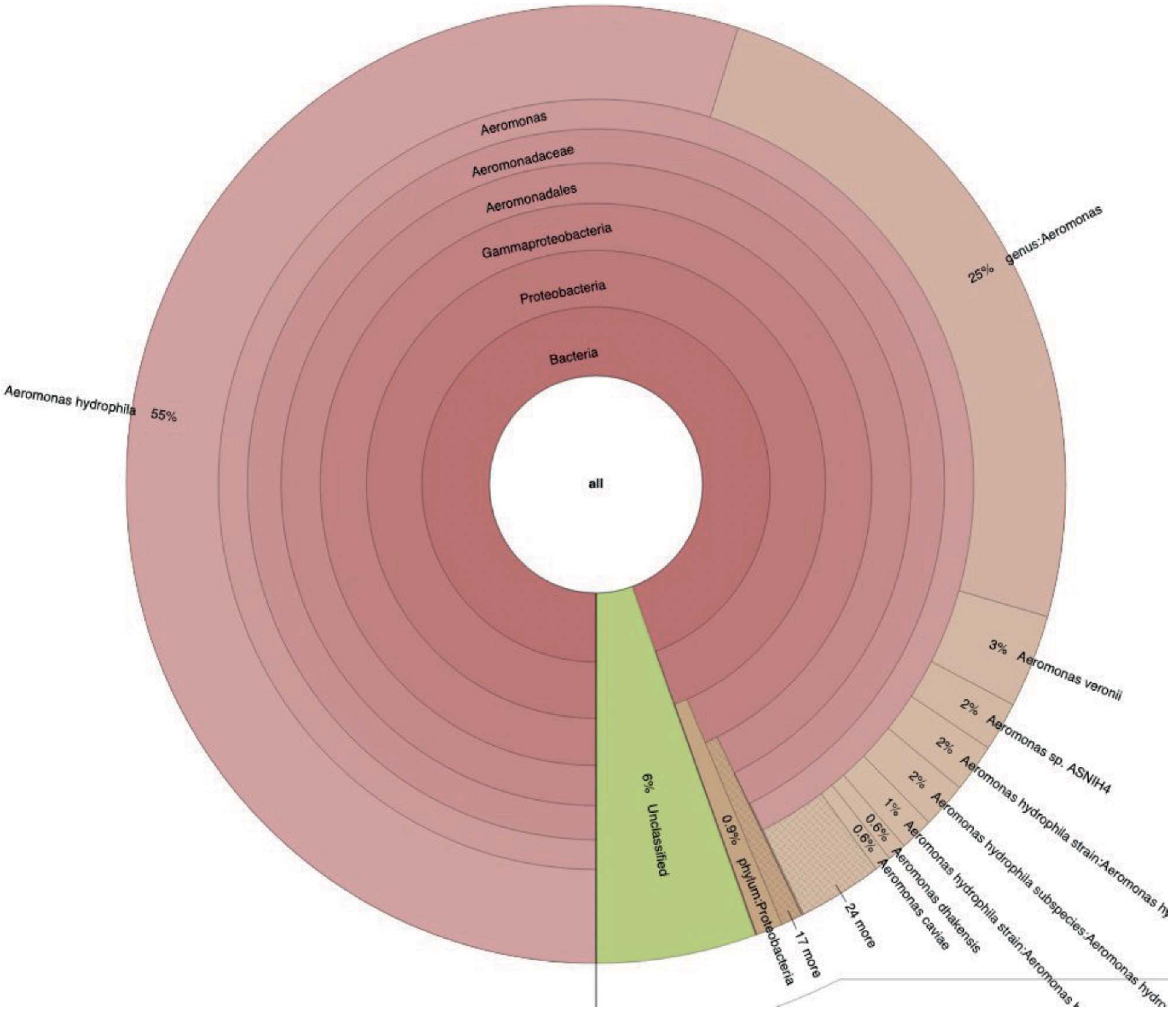
De novo sequencing and taxonomic classification show that an Aeromonas genus taxon can explain the deltamethrin tolerance. The ‘Sequana’ procedure generated a sequence of 4.9 Mbases submitted to a Kraken2 taxonomic classification. This taxonomic representation shows that the isolated taxon belongs to the Aeromonas genus with a high sequence identity with *Aeromonas hydrophila*.

We then plated the enteric bacteria content from ATB-treated 33S, and isolated 5 colonies randomly selected that were individually tested in sensitive *A. coluzzii* mosquitoes. The mosquitoes (maintained on sugar without ATB) were allowed to feed on naïve blood-fed (control group) or on blood supplemented with each of the isolated bacterial colonies. 48 h after the blood meals, all mosquito groups, including the control, were exposed to the deltamethrin. We found that only the mosquito group fed on Colony N°3 (C3) phenocopied the deltamethrin tolerance previously observed in mosquitoes treated with ATB (Fig. 4B).

De novo complete sequencing of bacterial DNA from C3 was performed. 4.9 Mb were generated, and taxonomic classification showed that the isolated taxon belongs to the *Aeromonas* genus and shares more than 55% sequence identity with *Aeromonas hydrophila* (Fig 5). This suggests this is probably a new, unidentified Aeromonas hydrophila taxon.

### Overall P450 activity does not explain deltamethrin tolerance in mosquitoes supplemented with bacteria

We tested and measured whether overall P450 activity could account for the delmethrin-tolerant phenotype. We used mosquitoes treated or not with ATB, blood-fed by supplementation (or not for the control) with the isolated *Aeromonas* taxa, and we prepared microsomal fractions to measure P450 activity using BOMFC as a model substrate. The kinetic parameters Vmax and Km are reported in Table 2.

**Table 2.** Kinetic parameters (Km and Vmax) of cytochrome P450 activities measured with BOMFC substrate in mosquito microsomal preparations. Values represent means ± SEM from three biological replicates (n=3 per condition; missing data for replicate number three for A.c 33S + ATB − BM and A.c 33S + BM + C3 bacterial clone due to insufficient protein quantity)

| **Mosquito** | **Vmax (pmol/min)** | **Km (μM)** |
| --- | --- | --- |
| A.c 33S - ATB | 0.454 ± 0.106 | 2.721 ± 1.415 |
| A.c 33S + ATB - BM | 0.424 ± 0.156 | 6.929 ± 2.545 |
| A.c 33S - ATB + BM | 0.177 ± 0.072 | 4.210 ± 3.013 |
| A.c 33S + BM + C1 bacterial clone | 0.216 ± 0.033 | 3.715 ± 1.912 |
| A.c 33S + BM + C3 bacterial clone | 0.192 ± 0.016 | 12.190 ± 0.988 |

No significant difference in reaction rate (Vmax) was observed between conditions (p=0.1294, one-way ANOVA statistical analysis), whereas significant variations were observed for Km (p=0.0257). Indeed, Km was higher for mosquitoes supplemented with bacterial clone C3 compared to other conditions (p<0.05, Turkey post-hoc tests), except condition A.c 33S + ATB - BM. Determining the kinetic parameters allowed us to define a dose close to Km in order to determine the specific P450 activity for all conditions (Fig. 6). No significant differences were observed between conditions.

**Figure 6:**
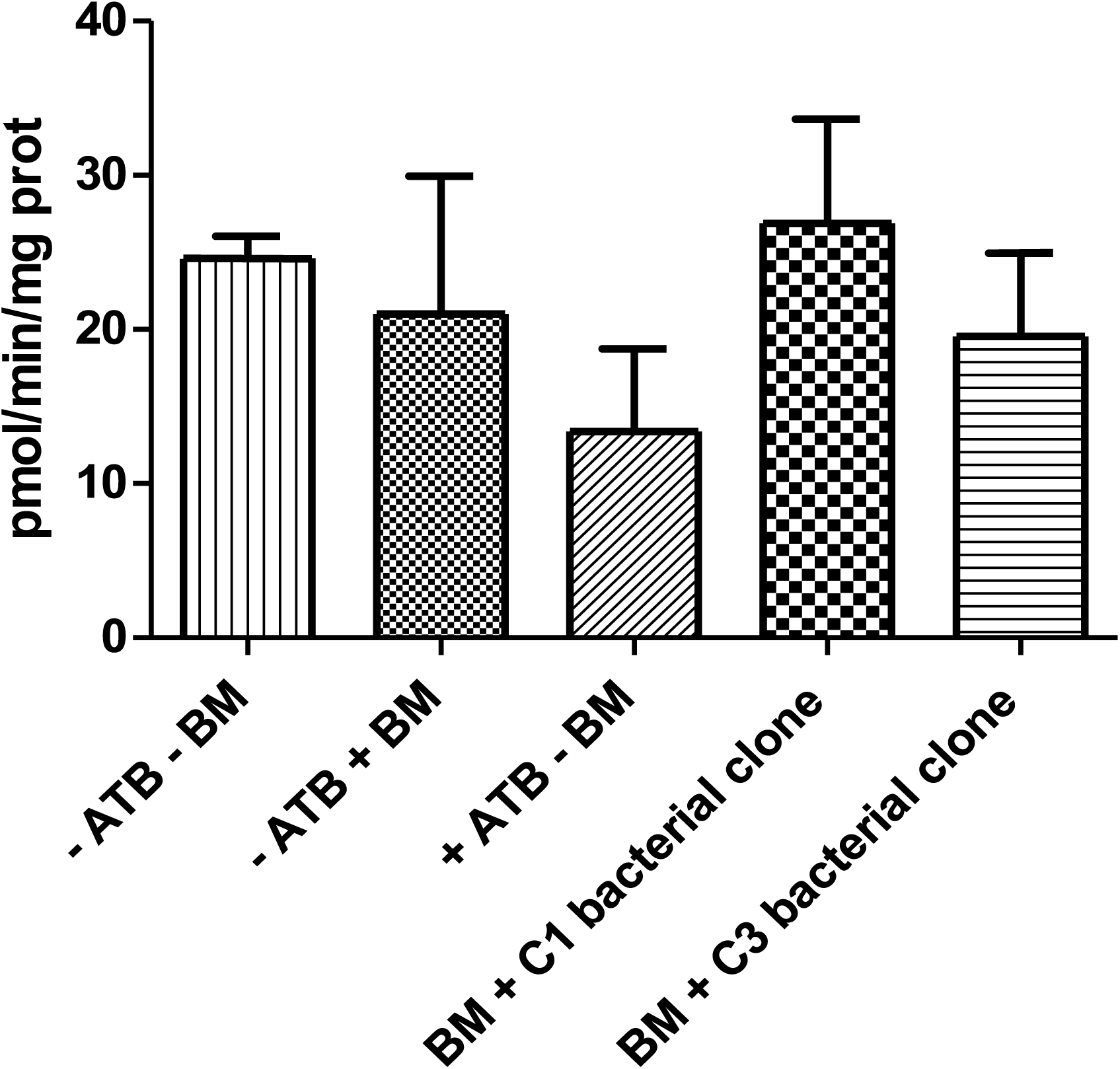
Specific cytochrome P450 activity measured with the model substrate, 7-benzyloxymethoxy-4-trifluoromethyl coumarin (BOMFC), at a dose close to Km. Data represent mean values of three biological replicates. Activities were statistically analyzed by one-way ANOVA and tukey test. -ATB= without antibiotics; +ATB= with antibiotics; -BM= no blood-meal; +BM= with blood-meal.

## Discussion

Unlike insecticide-resistant mosquitoes, the influence of gut dysbiosis triggered by antibiotic uptake in insecticide-susceptible mosquitoes has been poorly investigated. In the present study, we investigated the role of gut microbiome dysbiosis in insecticide susceptibility of malaria mosquito vector, *Anopheles*. We showed that antibiotic-mediated depletion of the enteric bacteriome on both sugar-fed and blood-fed mosquitoes confers deltamethrin tolerance to susceptible mosquitoes. This effect was phenocopied in different Anopheles mosquito colonies from distinct geographic areas and maintained in independent insectaries. This suggests that the underlying mechanism mediating deltamethrin tolerance under gut dysbiosis conditions is not strain-specific and is probably conserved. We found and isolated an ATB-tolerant bacterial taxon identified as an *Aeromonas*-like taxon sharing 55% sequence identity with *Aeromonas hydrophila*. The *Aeromonas* genus and particularly *A. hydrophila* have been identified in wild mosquitoes: in *A. gambiae* and *A. funestus* (44) and in Asian *Anopheles* (45), strongly suggesting that this taxon is not a lab artefact. In addition, antibiotic multi-resistant *Aeromonas*.spp, including resistant *A. hydrophila*, were isolated from different parts of the world and were reported to be mainly resistant to penicillin and ampicillin (46–49). *Aeromonas. sp* are common aquatic microorganisms found in irrigation or river water (50–53). Therefore, *Aeromonas*. sp could be acquired or maintained during the aquatic larval development of some mosquito species. Nevertheless, we could not exclude a maintenance of this *Aeromonas* taxon through vertical transmission as highlighted in other mosquito species (54).

Regardless of our findings, one hypothesis could be that the isolated ATB-tolerant Aeromonas taxon may undergo a penalized growth in normal sugar background by resource competition with other dominant taxa, whereas in ATB-treated background where resource competitions are reduced, it becomes dominant. *A. hydrophyla* is considered an important pest in aquaculture as it causes diseases called Motile Aeromonas Septicemia (55, 56). Interestingly, Nile tilapia fish infected with *A. hydrophyla* displayed an increased expression of P450 in their liver tissues (57).

Thus, one hypothesis is that the isolated *Aeromonas* taxon may secrete a factor that activates a detoxifying pathway such as the cytochrome P450, which can incidentally detoxify deltamethrin, leading to insecticide tolerance. Our current study investigated the mechanism that underlies this insecticide tolerance mediated by the ATB treatment, focusing on the metabolic detoxifying pathway involving P450 activity. Alteration of P450 activity either via PBO exposure or via cytochrome reductase (CPR) silencing, abolished or reduced deltamethrin tolerance triggered by ATB exposure. The high mortality found in the dsGFP control group compared to non-injected A.c 33S treated with ATB could be attributed to the injection event, where the cuticle property at the injection site would be modified and thus trigger a higher penetration efficacy of the insecticide. Nevertheless, the significant rescue of deltamethrin susceptibility in CPR silencing, together with the PBO phenotype, indicates that the activity of the P450s constitutes a mechanistic interplay for the insecticide tolerance mediated by the microbiome depletion.

We also investigated the expression of 7 key insecticide-resistance-associated P450s and GSTE2 by TaqMan qPCR in antibiotic-treated and non-treated mosquitoes. However, none of the tested genes was significantly modulated in their expression by the ATB exposure. Several hypotheses could be made in link with the detoxification process after ATB treatment: i) since the mosquito genome encodes ∼110 CYP450s (58), other P450s than the tested ones could be involved, especially since the tested P450s have been characterized in resistant *Anopheles* field mosquitoes, whereas in the current work, our phenotype is observed in susceptible lines, ii) a combinatorial activity of multiple P450s could explain the detoxification process, ii) depletion of the microbiome do not modify the expression but may have post-transcriptional influence on P450 and thus on their activity. For example, in mammals, the enteric microbiome increases the activity of certain P450 without modulating their expression (59). However, further works are required to test these hypotheses involved in this interplay with the microbiome.

In the current work, we also simulated mosquito blood meals on antibiotic-treated persons as they would occur in natural settings. Amoxicillin is a frequently prescribed antibiotic to treat respiratory infections, infections of the ear, nose and throat, as well as urinary tract and skin infections (35–37). Following a blood meal supplemented with Amoxicillin (0.2mg/ml), deltamethrin tolerance was recorded at 48 and 72 hours post-blood meal, whereas the control group fed on untreated blood displayed 100% mortality. The lack of phenotype at 24h post-feeding could be assigned to the proliferation of the bacteria gut flora after the blood meal, which reached a pic at 24h even with the addition of the ATB as recorded by the 16S level, and consistent with the data found in (60). In addition, this slight delay observed in the phenotype also suggests that the influence of the microbiome reduction in insecticide susceptibility is indirect and may requires the activity of the P450s as an intermediary. Deltamethrin tolerance caused by antibiotic exposure of susceptible mosquitoes may impact residual transmission in African settings. Further field works are required to assess whether the high prevalence of antibiotic administration or self-medication prevalence in specific areas could influence insecticide efficacy and thus the vectorial capacity. If this is the case, impregnated bed nets combined with deltamethrin and an efficient antibiotic against *Aeromonas. sp* and formulated to penetrate mosquito cuticle could be used to reduce the deltamethrin tolerance and thus vectorial capacity.

**Figure S1: Antibiotic efficiency**

**A.** 16S qPCR on DNA samples from Penicillin/streptomycin and Gentamycin-treated mosquitoes show a decreased abundance of the enteric bacteriome. **B.** Mosquitoes fed on blood supplemented with amoxicillin display decreased abundance of the enteric bacteriome measured via the 16S level by qPCR. The 16S decrease reaches a maximum at 48 hours post-blood feeding.

**Figure S2: The 16S decrease in abundance explains the deltamethrin tolerance effect**

**A.** Antibiotic-treated A.c 33S survivors from a first insecticide exposure were returned to normal sugar meals (without antibiotic) for 48 h and directly re-exposed to deltamethrin (0.05%). Stopping the ATB pressure for 48 h reverted A.c 33S to their original insecticide susceptible status (∼97% mortality). **B**. Tetracycline-treated mosquitoes triggered the same tolerance phenotype in mosquitoes exposed to deltamethrin compared to the untreated control group. **C**. Tetracycline efficiency on the enteric bacteriome abundance was measured by qPCR quantification of the 16S level in antibiotic-treated versus untreated mosquitoes (dotted line). The results show a 50% decrease in the abundance of the 16S. *=p<0.05

**Figure S3: Silencing efficiency of CYP6M2 and CPR**

**A.** The histogram shows the silencing efficiency of the CPR gene in mosquitoes injected with dsCPR compared to the control group injected with dsGFP (dotted line). The results show a decrease expression of the CPR gene at 60%. **B**. The histogram shows the silencing efficiency of the CYP6M2 gene in mosquitoes injected with dsCYP6M2 compared to the control group injected with dsGFP (dotted line). The results show a decrease in the expression of the CPR gene at 70%. *=p<0.05; **=p<0.01

## Acknowledgements

We thank the Institut Pasteur core facility, the Center for the Production and Infection of Anopheles (CEPIA) for rearing of mosquitoes.

## Funding

This work received financial support from the European Commission, Horizon 2020 Infrastructures #731060 Infravec2; European Research Council, Support for frontier research; and French Laboratoire d’Excellence “Integrative Biology of Emerging Infectious Diseases” #ANR-10-LABX-62-IBEID.

## Notes

### Competing Interest Statement

The authors have declared no competing interest.

