## Supplementary figures and images for "Anopheles resistance to deltamethrin can be caused by the increased abundance of an enteric Aeromonas taxon"

### Supplemental Figure 1

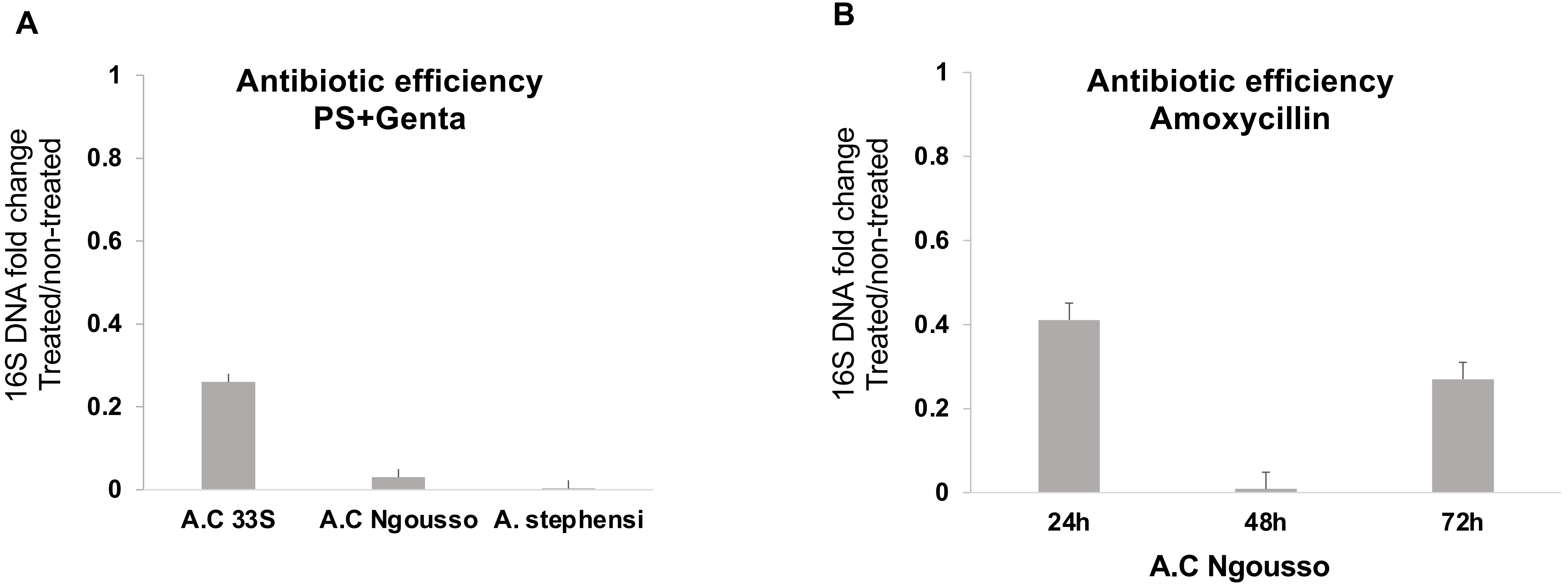

### Supplemental Figure 2

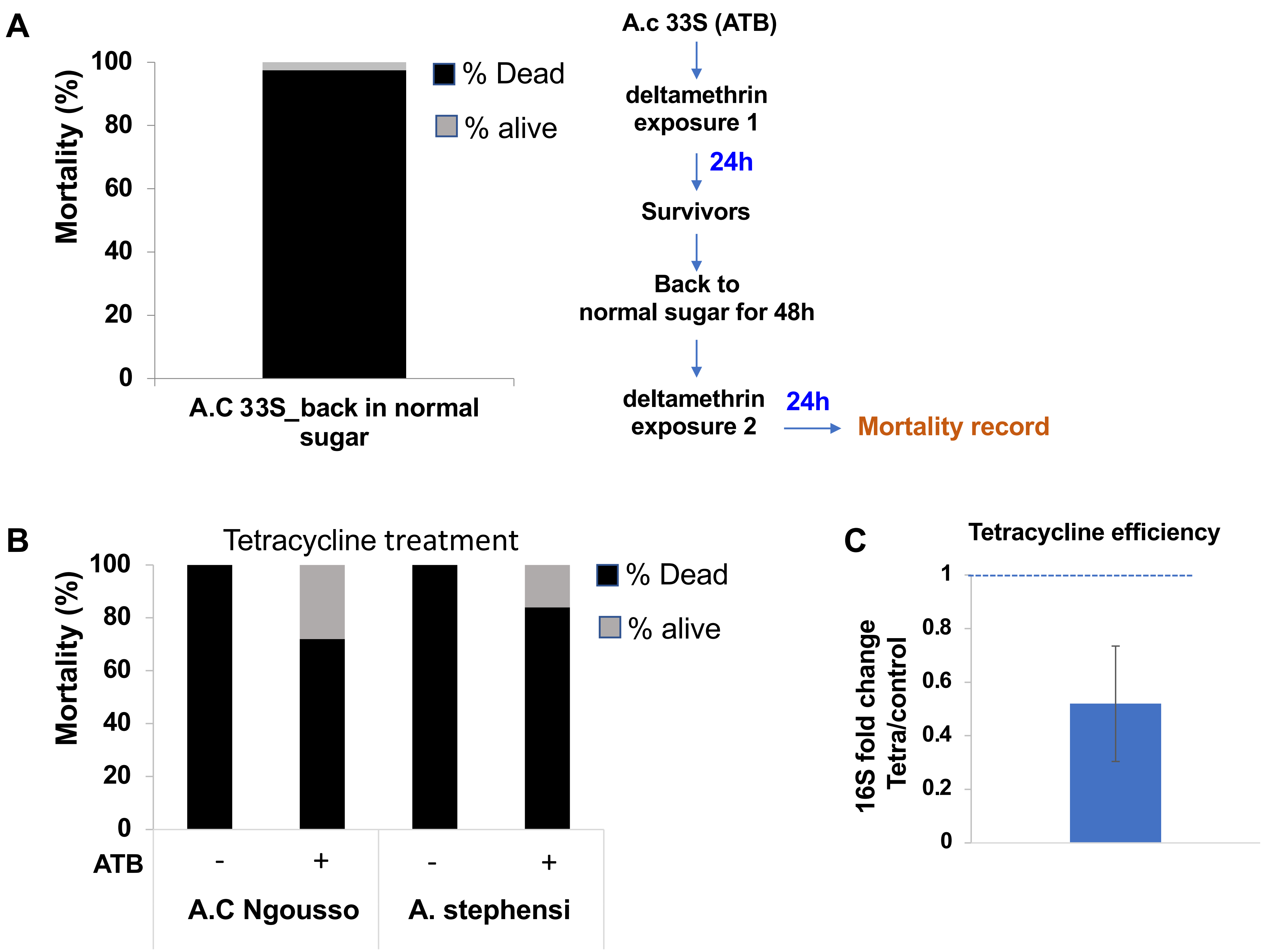

### Supplemental Figure 3

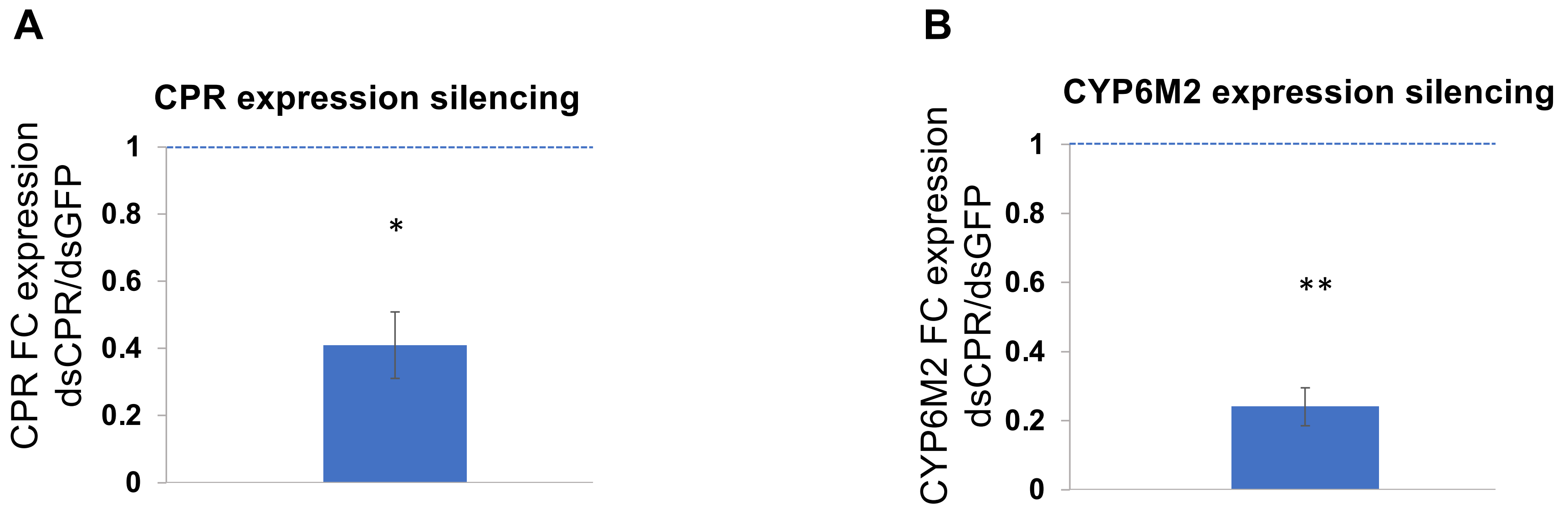
